# Genetic Engineering with Quantum Circuits: creating codes and studying BioBloQu genetic elements

**DOI:** 10.1101/2025.05.02.651535

**Authors:** Patrícia Verdugo Pascoal, Deborah Bambil, Rayane N. Lima, Marco Antônio de Oliveira, Luisa M. A. de Tacca, Mendeli H. Vainstein, Elibio Rech

## Abstract

Quantum biology is an emergent field that investigates quantum-mechanical phenomena, such as superposition, tunneling, and entanglement, in the context of data manipulation from living systems. The exploration and engineering of nucleotide sequences rely on quantum mechanical principles, particularly the use of qubit states for the development of quantum codes. Biological sequencing data is produced at about 1 Gb/h, but analysis lags due to complexity and the limitations of classical computing. Despite these challenges, quantum computing offers a potential tool for analyzing and assembling biological data. Here, we developed quantum codes for genetic engineering. The developed quantum computational framework identifies sequences of interest within a genomic database. It locates the left and right boundaries of the scar region in the JCVI-Syn3B genome and detects 20 nucleotides flanking each boundary. After confirming the left and right ends of the scar, a secondary computational routine performs the targeted insertion of the BioBloQu structure, composed of genetic elements, into the previously characterized scar region. Our algorithms constitute a unique starting point for advanced genetic data manipulation. This tool could accelerate the exploration of large volumes of genetic data and enable the developmental design and assembly of synthetic genomes enhanced by quantum computing. Further improvements in algorithms and codes, along with the expanded availability of devices, will accelerate the search for data for applied genetic research.

---

The intersection of quantum theory and biology has intrigued researchers for over a century^1^, suggesting a fundamental limit to our understanding of life through the lens of atomic structures. Following Bohr’s formulation of the atomic model in 1913, physics sought a more consistent theoretical foundation for subatomic phenomena. In 1925, Werner Heisenberg developed a new approach based on observable quantities, introducing the formalism of matrix mechanics. With the support of Max Born and Pascual Jordan, this theory was consolidated as the first coherent formulation of quantum mechanics. This breakthrough redefined the foundations of modern physics and earned Heisenberg the 1932 Nobel Prize in Physics, marking the beginning of contemporary quantum theory^2^. In 1996, Grover introduced a quantum algorithm that revolutionized database searches, reducing the complexity from O(N) to O(√N); however, its practical applications remain limited by the operational demands of large databases^3^. The advent of the idea of building quantum mechanical computers in the early 1980s ^4^ set the stage for groundbreaking advances, culminating in the late 1980s and early 1990s with pioneering descriptions ^5-8^ aimed at solving problems previously considered intractable with classic algorithms.

In the coming years, quantum simulations are expected to make significant advances in areas such as quantum chemistry and high-temperature superconductivity. Recent studies indicate that, despite notable advances in quantum computing, the realization of fault-tolerant, fully scalable quantum systems suitable for broad practical applications is expected within the next two decades. The current challenges include protecting the delicate states of quantum bits (qubits) from errors and unwanted environmental disturbances. This gap between the theoretical potential and the current technological capabilities underscores the transformative promise of quantum computing, particularly in biological research ^9^. Realistic expectations for advances in quantum computing are needed, large companies claim to offer quantum computing services while remaining limited to small-scale algorithms ^9^.

Quantum computers operate using quantum bits (qubits), which, unlike classic bits, can exist in multiple states simultaneously because of superposition. Quantum entanglement establishes correlations between qubits, providing a fundamental feature that allows for gains in computational speed in quantum algorithms. ^10-12^. Importantly, quantum computers should not be seen solely as an evolutionary successor to classical systems ^13^; instead, they excel at solving specific, complex problems, whereas classical systems remain optimal for other tasks ^14^.

Quantum computing opens new avenues for biological research that could lead to breakthroughs in medicine ^15^, biotechnology ^16,17^, and our understanding of life itself ^18^ and evolution ^19^. Currently, quantum computing approaches, with their unique capabilities, offer promising solutions for the acceleration, prospection and manipulation of nucleotide sequences with the aim of discovering specific and innovative traits in genomics ^20-24^. For a broader perspective on quantum computing in biology, several articles have been published ^16,25-34^. However, the lack of sufficient quantum algorithms and codes tailored to computational biology challenges hinders the potential of quantum computing to accelerate genetic data processing and manipulation in this field ^10,11,13,14,35-45^. New formulations for RNA structures were introduced and reported using quantum pattern recognition and the quantum Hamming distance within Grover’s framework, enabling the extraction of linear regions from complex forms ^38^. To maximize the achievable quantum speedup, methods have been developed for concatenating different linearized segments of RNA structures into longer sequences. The mechanism was simulated using IBM’s Qiskit quantum simulator by employing RNA sequences from GenBank and mirBase ^38^. Iteration-complexity comparisons demonstrated the potential for quadratic speedups in quantum computing over the classic methods, and simulation results indicated high search accuracy. A quantum computing method to improve the process of assembling DNA sequences was developed. The approach combines data from two types of sequencing technologies, one that produces short pieces of DNA and another that produces longer ones, to build more complete and accurate genetic maps. Using a divide-and-conquer strategy, the method reduces the amount of quantum resources needed, making it practical for current quantum computers. The algorithm also benefits from quantum effects, such as entanglement, which help it find better solutions faster^46^. Using quantum channels and quantum walks, a quantum approach to simulating evolutionary relationships among species reproduced classical phylogenetic models and enabled likelihood estimation from genetic data. While the method matches classical outcomes, it also points to future use of quantum coherence for deeper evolutionary insights. The work establishes a foundation for applying quantum computing to phylogenetics^47^.

These results suggest that future quantum search methods could significantly accelerate the genome sequencing process as quantum computers become more widely adopted. Further research should investigate the effects of decoherence on detection accuracy and assess the computational demands of each iteration on actual quantum hardware.

A quantum search using Grover’s algorithm showed the explicit construction of a gate, programmed in Qibo for N = 8 and M = 2. The database size was set to N = 8 possible entries, among which M = 2 entries were defined as marked targets (i.e., the desired search results). This setup demonstrates how quantum parallelism can effectively reduce the computational complexity associated with locating specific items within an unstructured dataset. While the classic algorithm required N + M queries in the worst case, the quantum algorithm found the solution with only one or two queries, demonstrating an improvement in execution time. However, a limitation of the algorithm is that N must be significantly greater than M. In addition, a four-dimensional extension of the algorithm was implemented to perform codon searches in DNA sequences using ququarts (four-dimensional qudits), with an adapted alphabet containing A, C, G, and U ^38^. In this context, advancing novel quantum algorithmic methodologies is a central challenge for the evolution of quantum information science. Addressing this challenge necessitates deep interdisciplinary collaboration between researchers in quantum physics and computational biology, uniting theoretical foundations with biological data modeling ^27^.

We developed and implemented two codes that apply quantum algorithms to optimize large-scale genetic sequence analyses using 27 qubits derived from the IBM simulator. These algorithms initially perform a search for a specific nucleotide sequence (e.g., the sequence pattern of a gene of interest) within a nucleotide database (e.g., an assembled genome, metagenome, or similar dataset). Subsequently, a second algorithm identifies the scar region within the minimal genome model (JCVI-Syn3B), accurately defining and detecting the 20-nucleotide boundaries at both the 5′ and 3′ ends. The scar region is a residual sequence of nucleotides that remains in the genome after a gene-editing process, such as the insertion or deletion of an element. Unlike a simple deletion, a scar is a specific, intentional byproduct of site-specific recombination, serving as a permanent record of the operation. Its primary function is to serve as a unique and unambiguous marker that identifies and/or confirms the desired editing event. In minimal genome models, the automated recognition of these scar regions is crucial for tracking and validating the multiple modifications performed, ensuring the integrity and accuracy of the constructed synthetic genome. Following the identification of the gene sequence of interest and the detection of the scar region in the minimal genome, a secondary quantum algorithm constructs and integrates a functional genetic element, designated BioBloQu, directly into the recognized scar site. This insertion process simulates a modular synthetic biology operation, enabling programmable genome editing and high-fidelity *in silico* modeling of synthetic biological constructs by applying quantum computing principles.

This study proposes an extension of a quantum search framework tailored for big data applications. The system serves as a computational platform enabling the design, construction, and hierarchical assembly of functional biological building blocks for synthetic and complex genome engineering.

## Results

### Searching for the sequence of interest using the QuBio.py code

The primary objective of the quantum simulation conducted with *Qubio*.*py* was to evaluate the algorithm’s effectiveness at identifying specific DNA (target sequences) of 50 nucleotides within a genetic database (3022 nucleotides). The target DNA sequence was selected based on its relevance to ongoing gene identification research; in this case, a conserved functional domain of the Cas9-like nucleotide sequence found in Brazilian biomes was employed.

This quantum code performs nucleotide sequence comparisons, allowing a mismatch rate of up to 30%. This approach differs from BLAST ^48^, a widely utilized tool for sequence alignment that, although efficient, is predominantly classical. It allows up to 30% differences between the compared sequences, which can be useful in contexts where sequence variations are expected, such as in evolutionary studies or analyses of homologous genes. Although BLASTN also identifies similar sequences, its calculation similarity values and handling of mismatches is more complex, accounting for factors such as gap penalties and different forms of similarity.

Focused on applications where speed and tolerance to variation are paramount, the method can be particularly useful for projects that require rapid sequence analyses with a focus on general similarities.

Our code represents an interesting advance in the application of quantum computing to sequence comparisons, enabling fast and flexible analyses in contexts where tolerance to mismatches is desirable. This innovative approach can complement the classic tools, offering new perspectives for genomic research.

The simulation and implementation stages were performed on the IBM Quantum Experience platform, which used a quantum processor with 27 qubits^49^. The Grover’s algorithm-based code was configured with a search space corresponding to a genetic database sequence of 3022 nucleotides from a Brazilian biome metagenome database^50^. On the IBM simulator, the algorithm completed the search in 1 second, with 10 iterations. This difference was expected due to the additional overhead and complexity of executing the algorithm on physical quantum hardware compared to a simulated environment. Overall, the real quantum computer was slightly slower but still efficiently handled the same sequence size and number of shots as the simulator, indicating its practical applicability for such bioinformatics tasks. The results of the tests are summarized in Table 1.

**Table 1.** Comparative performance of different computers in terms of executing nucleic acid sequence search algorithms.

|  | Classical<br>computer/BLASTN* | IBM Simulator |
| --- | --- | --- |
| Time of running (seconds) | 0.342 | 1 |
| Iterations | N/A | 10 |
| Target_sequence size (nt) | 50 | 50 |
| Database_sequence (nt) | 3022 | 3022 |

\*128 CPU Processor.

Additionally, by performing 10 iterations of the test (Fig. 2), it was possible to verify an oscillatory pattern in the final probability of the sequence being found in the database. The probability of identifying the best candidate changes during the search process, using *K = 20* possible candidates, which corresponds to the 20 different starting positions in the database where the target sequence could potentially match. The x-axis indicates the number of Grover iterations, while the y-axis shows the probability of obtaining the correct candidate at each step. The curve follows the typical sinusoidal pattern of Grover’s amplification (Fig. 2). As the number of iterations increases, the probability of finding the correct solution rises until it reaches a peak (around iteration 4, with a probability close to 1.0). After this point, the probability begins to decrease, reflecting the periodic nature of quantum interference in the search space. This confirms that the pseudo-Grover model accurately captures the expected quantum behavior. The number of iterations can be understood as the depth of the quantum search through biological data. The peak at iteration 4 marks the optimal point where the system maximizes the likelihood of identifying the correct target sequence before the probability begins to oscillate again.

**Fig. 1.**
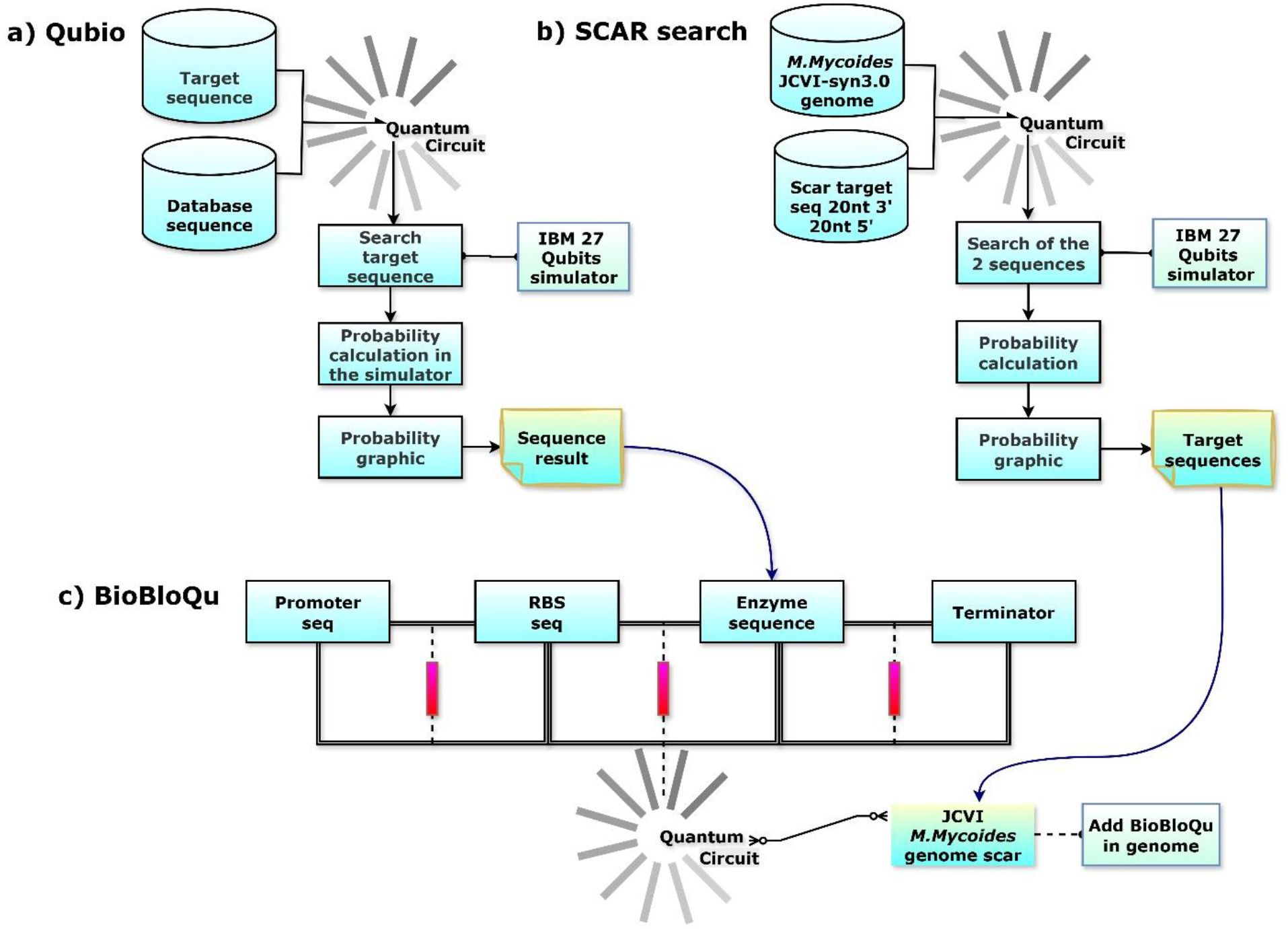
Workflows for genome analysis and synthetic assembly with a quantum computing approach. **a - Qubio**: Simulation of a quantum-inspired search algorithm applied to genetic sequences. A target sequence and a database sequence are encoded as qubit states within a quantum circuit. Probability amplitudes are computed via Grover-type iterations to identify the most probable matches, illustrating how quantum search can accelerate pattern recognition in genomic data. **b - SCAR search**: Quantum simulation of scar-sequence detection within the *Mycoplasma mycoides* JCVI-Syn3B genome. The algorithm identifies genomic scar regions-standardized recombination or insertion sites-based on probability distributions generated by the IBM 27-qubit simulator. These loci represent potential manipulation points for genome editing. **c - BioBloQu**: Quantum circuit-assisted assembly of functional genetic modules (promoter, ribosome-binding site, enzyme gene, and terminator) designed for insertion into the identified scar regions of the *M. mycoides* genome.

**Fig. 2.**
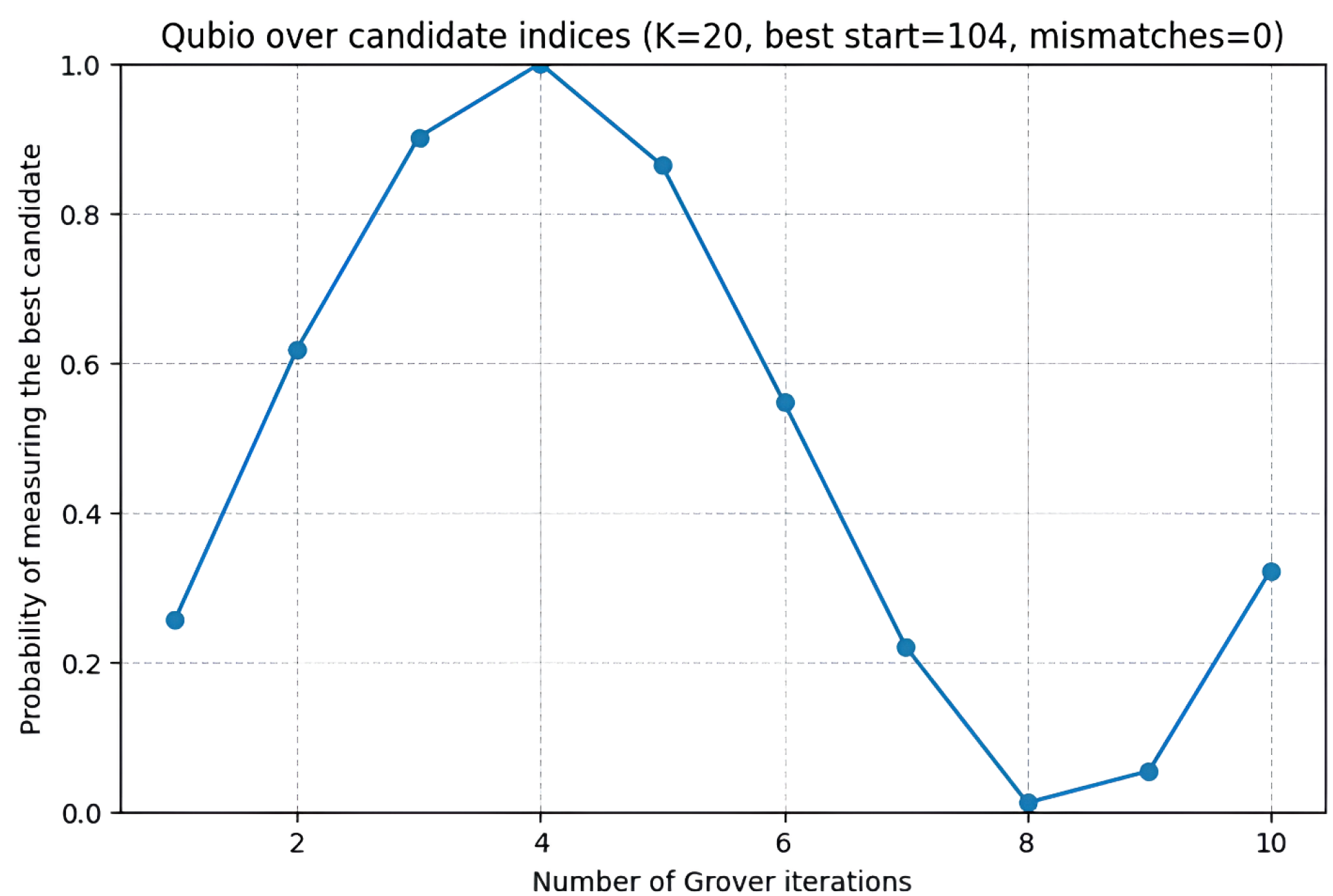
Probability of finding the target sequence in the database of Brazilian metagenome sequences according to the QuBio.py code. Running time: 10 iterations ran in 1 second using 27 qubits of the IBM simulator.

We also tested how the code behaved by changing only 15 nucleotides in the target sequence (30% mismatch rate) while keeping the database unchanged. The results indicate that the algorithm was sensitive to this specific change. When we changed to more than 15 nucleotides, we observed a sequence-finding probability of zero.

Additionally, we performed a predictive analysis using a linear regression model to compare the predicted processing times using a quantum computer with various qubit counts (150, 200, 250, and 300 qubits) and a traditional server with 100 threads. These predictions were made for target sequences of 100, 200, and 300 base pairs, using the actual processing times obtained from a 100-qubit quantum computer and a 100-thread server, with 50 base pairs as a reference. The search was conducted on a 1-petabyte database containing a 3-Kb reference sequence. The results indicate that, as expected, increasing the number of qubits significantly reduced the processing time. For example, the overall trend reveals a direct relationship between sequence length and computational time for all configurations. However, increasing the number of qubits leads to a substantial improvement in performance and scalability. The system configured with 100 qubits exhibits the highest processing times, rising from approximately 36 minutes for 50 base pairs to around 122 minutes for 300 base pairs. With 150 qubits, the time decreases slightly, ranging from about 34 to 100 minutes across the same interval. The 200-qubit setup shows a more significant reduction, processing from roughly 32 minutes at 50 base pairs to about 80 minutes at 300 base pairs, representing an improvement of nearly 30% compared to the 100-qubit configuration (Fig. 3).

**Fig. 3.**
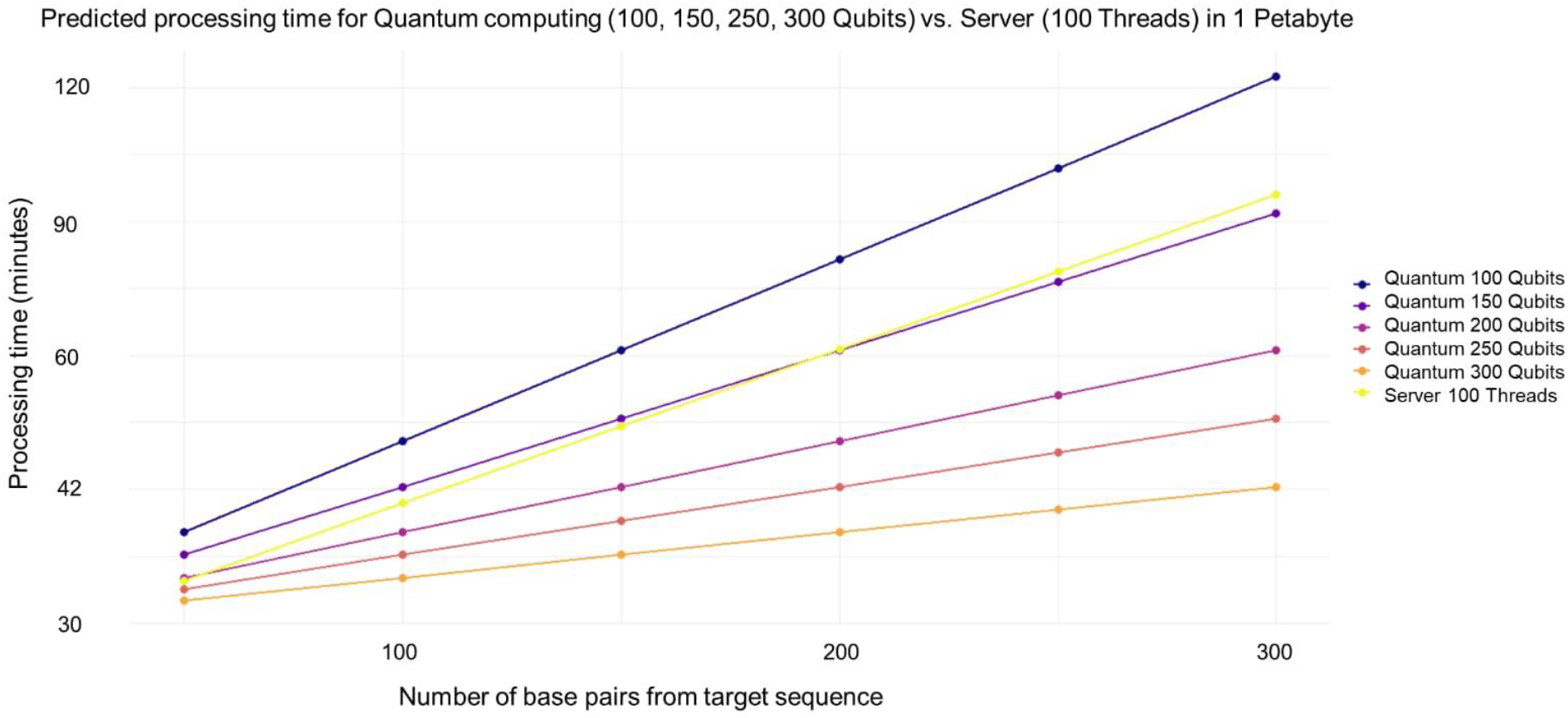
Estimated processing times for sequence analysis using classical and quantum computer approaches. Predicted processing times obtained from a linear regression model comparing simulated quantum computing performance (assuming 100– 300 qubits) with a traditional high-performance server using 100 threads. The estimates correspond to target sequences of 100, 200, and 300 base pairs analyzed within a 1 petabyte dataset. These results represent theoretical projections rather than measurements from an existing quantum device, providing an indicative comparison of expected computational scaling between quantum and classical approaches.

These results suggest that quantum computing offers substantial improvements in processing efficiency, particularly as sequence length increases, making it a promising solution for handling large-scale metagenomic data.

### Finding a scar in the minimal M. mycoides JCVI-Syn3B cell and building functional biological blocks

New genes identified in metagenome studies, as exemplified above with Cas9 from Brazilian biomes^50^, can be sources of new functions and attributes for genetically modified or even synthetic organisms. Therefore, quantum-computing-based tools that assemble functional genetic elements and detect optimal insertion sites will be valuable techniques for guiding rational genome editing and even the *de novo* design of completely synthetic genomes. In that context, we developed a second algorithm to execute these tasks.

We worked with sequences acquired from the minimized *M. mycoides* JCVI-Syn3B genome, given its importance as one of the prominent landmarks in synthetic biology. The first breakthrough in the development of this minimal cell was the creation of *M. mycoides* JCVI-Syn1.0, the first cell controlled by a genome completely chemically synthesized in the laboratory^51^. This strain then served as a chassis for studies in the identification of essential genes and the removal of nonessential genes to generate a minimized version of the original genome, resulting in *M. mycoides* JCVI-Syn3.0, which included less than 50% of its parental genome^18^. Further iterations, including reintroduction of 19 previously deleted non-essential genes for morphological stability and addition of a landing pad region resulted in the derived strains JCVI-syn3A e JCVI-syn3B^52^.

Our second code was first used to identify the unique sequence present in Syn-3B and resulting from the deletion of the nonessential gene *mmsyn1_0531*, which was initially present between the essential genes *mmsyn1_0530* and *rrl* in Syn-1.0. This new gene organization scheme generated a new sequence that can be seen as a scar marking the deletion site of a nonessential gene. Each bar represents the probability of identifying the insertion site (or scar) for a specific target sequence: “AAAATCTGTCATAAATTATC” and “ATTATTCTCCTTTCTTTAGT”. The red dashed line indicates the theoretical maximum probability (1.0), corresponding to perfect sequence detection. Both target sequences exhibited probabilities around 0.5, suggesting that the circuit successfully identified the correct genomic regions but with residual uncertainty due to the limited number of Grover iterations and the size of the search space. These results demonstrate consistent quantum search efficiency in detecting scar regions for subsequent biological block (*BioBloQu*) insertion (Fig. 4). Genome assembly markers, commonly called scars, represent DNA sequences manipulated during editing, synthesis, construction, and genome engineering^53^. This region could be an interesting site for exogenous gene insertion, as it poses no risk of disrupting existing transcriptional units. Additionally, given their originally distant positions, suggesting independent regulation for each gene, a reduced degree of structural interference with gene expression was expected.

**Fig. 4.**
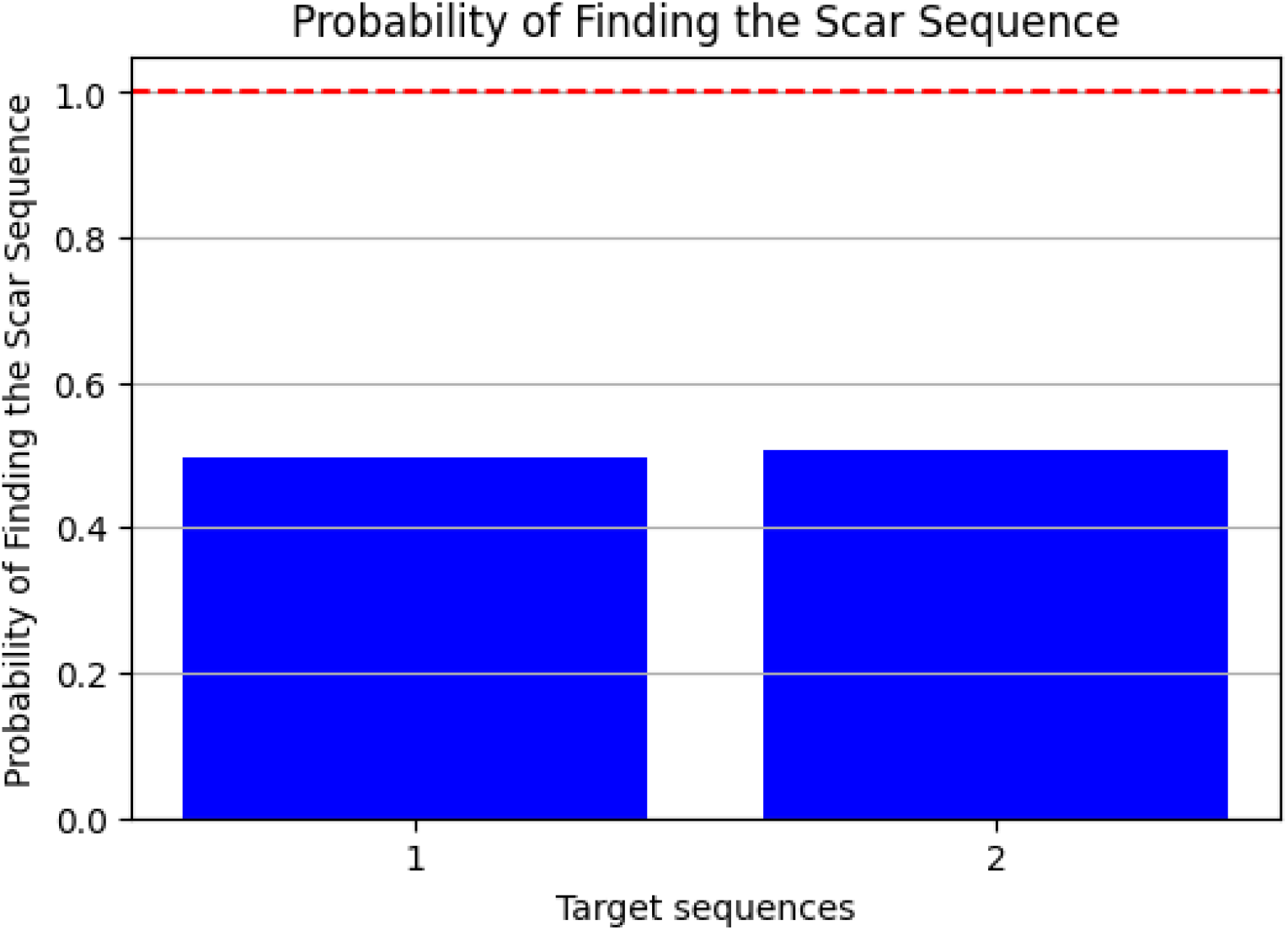
Probability of finding the scar sequence using a search algorithm in a quantum-based BioBloQu assembly model. The figure illustrates the probabilities of locating two target sequences within the reference genome after execution of the quantum circuit simulated on the Qiskit Aer Simulator. Each bar represents the probability of identifying the insertion site (or scar) for a specific target sequence.

Following the identification of an insertion site, the code then assembled a genetic structure with key components to create a transcriptional unit (a promoter, an RBS, a protein sequence, and a terminator). The assembled structure was named a BioBloQu (Fig. 5).

**Fig. 5.**
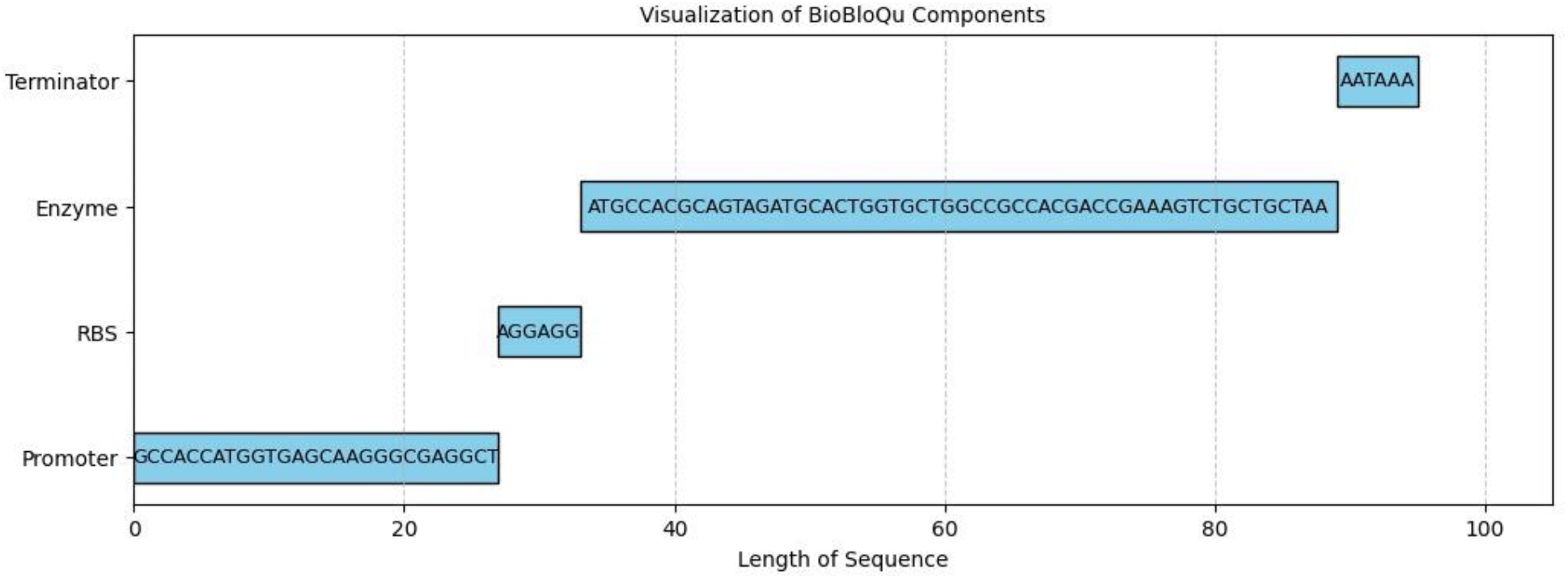
Representation of the structures of the components contained in a BioBloQu.

The generation of biological blocks for the construction process, which we denote as the BioBloQu of a synthetic sequence of interest using the search performed in step 1 with QuBio.py, included the addition of a promoter sequence, an RBS, the sequence of the enzyme of interest, and a terminator (which was able to add repressor and enhancer sequences).

This is a visual representation of the power of quantum computing for constructing genetic blocks inspired by BioBricks™. This approach not only streamlines the design process but also enhances the potential for creating innovative biological systems. As quantum technologies continue to advance, their applications in synthetic biology promise to revolutionize our ability to engineer life and tackle complex biological challenges.

This new genetic structure replaced the scar initially identified by the code, generating a modified minimal cell capable of secreting the protein presented in the designed BioBloQu (Fig. 6).

**Fig. 6.**
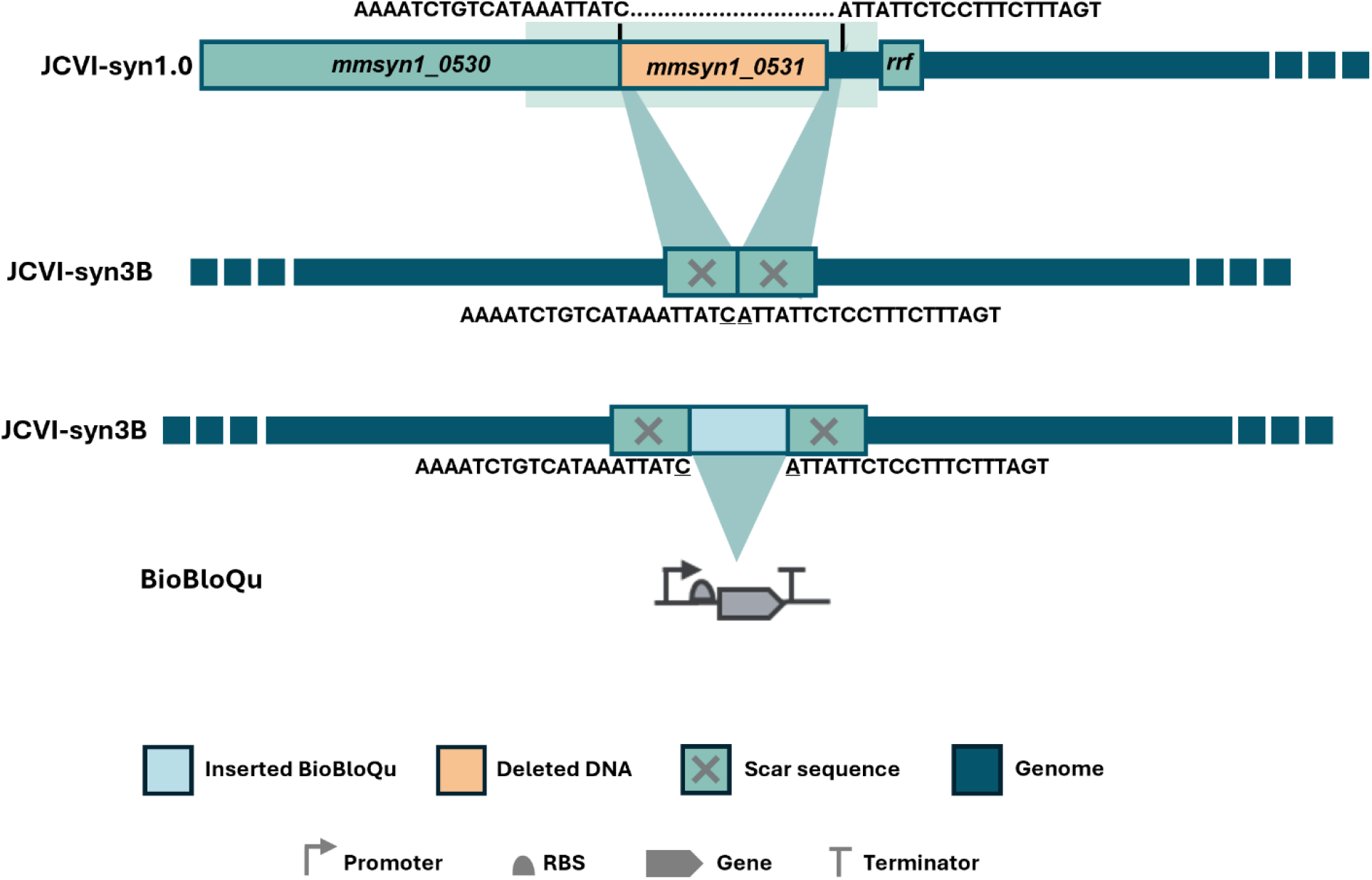
Integration of BioBloQu. The diagram illustrates the genetic logic and structural components involved in the targeted integration of the BioBloQu module. At the top, the native genomic locus (JCVI-Syn1.0) is shown, containing the *mmsyn1_0530* and *mmsyn1_0531* genes, as well as the *rrf* element. Flanking homology regions guide the recombination event. The central blocks marked with an “X” represent the scar sequences. After integration, BioBloQu is inserted between the scars, allowing the incorporation of additional functional elements, such as a gene cassette or a regulatory module.

The addition of these components suggests an effort to modify or enhance the genome’s functionality, possibly to engineer new phenotypic traits or optimize metabolic processes. In this way, quantum computing and the construction and optimization of genetic sequences are possible. This approach enables the exploration of computationally intensive genetic combinations using traditional methods, such as time-consuming bioprospecting, thereby highlighting the intersection of biotechnology with quantum computing. It serves as an example of how synthetic biology can be enhanced by emerging technologies, offering new possibilities for research and applications in diverse fields, such as agriculture and biofuels.

## Discussion

A quantum circuit framework was proposed in this study to estimate the universal distribution by simulating the superposition. This approach offers a quantum resource complexity level with linear data-size complexity, achieving a polynomial speedup over the classic methods. Thus, the exploration of program-output relationships represents a promising avenue for accelerating quantum searches, with specific data properties inferred from the universal distribution in the quantum superposition task ^29^. Some biomedical scientists are exploring how quantum computing can enhance algorithms and machine learning methods in various biological fields, such as protein design. As biologists adopt this technology, they seek to understand its underlying mechanisms and determine when classic computers are sufficient ^23^.

Several approaches have been proposed to optimize DNA sequence searches, from classic algorithms based on string comparisons to preliminary quantum implementations that seek to exploit quantum superposition and interference. However, gaps remain regarding the practical applications of these methods in large genomic databases.

From a biological standpoint, the quantum framework employed here successfully modeled the probabilistic oscillations associated with candidate selection, as evidenced by the amplitude peaks and periodicity observed in the Grover iteration simulations. These oscillations represent the interference dynamics that drive quantum optimization, where the probability of identifying the correct genomic or nucleotide target increases up to a maximum and then decreases with over-rotation in Hilbert space. The optimal point, observed around the fourth iteration, defines the balance between convergence speed and computational efficiency, illustrating how quantum interference can be leveraged for biological search optimization.

The mechanisms by which genetic information is encoded in nucleotide sequences were investigated through comparative analyses ^13^. Variations can occur at the nucleotide level or collectively due to recombination or deletion. Detecting these differences is essential for biology and medicine but requires substantial computational power due to genomic complexity. To address this issue, the authors proposed the use of flexible representations of quantum images (FRQIs) for sequence analysis purposes, enabling detailed comparisons and improving the detection accuracy achieved for subtle genetic variations^54^.

After analyzing comparative predictions of processing times using a quantum computer and our robust classical server with 100 threads (Fig. 3), the results suggest that quantum computing offers substantial improvements in processing efficiency, particularly as the sequence length increases, making it a promising solution for handling large-scale metagenomic data. Therefore, to match our server with 100 processing threads, the quantum computer needed approximately 150 available qubits. IBM’s quantum computer currently has 400 qubits, but fewer than 200 are available for use in either simulations or as real quantum units ^55^. Even at a smaller scale, quantum computing can offer efficiency gains as the number of base pairs in the target sequence increases. Using a larger number of qubits (e.g. 250 or 300) resulted in processing times that were significantly lower than those of classical computing, suggesting that quantum computing may be better suited to problems that require high computational power. The representation of classic computing shows a linear increase in processing time, indicating that as data increases, classic computing may become impractical for tasks that require intensive processing. These observations highlight the potential of quantum computing to transform fields that require the analysis of large amounts of data, such as bioinformatics, molecular modelling, and other scientific applications. Challenges remain, and there is a long way to go before quantum computing becomes established. We are working in parallel with this progress to maximize its capabilities and expand its achievements in biology and biotechnology.

Here, the use of 10 iterations is sufficient for the objective requires less time and computational power, as it takes approximately 1 second per iteration and 49 seconds for 100 iterations. Increasing the number of iterations in a quantum search algorithm can decrease the probability of finding the target, due to several factors inherent to quantum algorithms.

The potential of quantum computing to accelerate the discovery of specific and innovative traits from extensive nucleotide sequence data is immense. By providing unparalleled processing power, enhancing the precision of trait discovery, optimizing genetic algorithms, integrating with AI, and enabling real-time analyses, quantum computing could revolutionize how we approach genomics and trait development ^56^. As this technology matures, it promises to unlock new possibilities in biotechnology, agriculture, and medicine, paving the way for ground-breaking innovations that could reshape our understanding of life ^57,58^.

Algorithms can process large datasets more efficiently, reducing the time required for drug discovery processes ^15^. The use of quantum computing can lead to more accurate predictions of molecular properties, facilitating the identification of promising drug candidates. However, their paper also had several limitations. The current state of quantum computing hardware is still in its infancy, which limits the practical implementations of quantum machine learning algorithms in real-world drug discovery scenarios.

The study published by Rodríguez ^38^, which also used the approach in our code, exemplified the trade-offs between computational complexity and algorithmic effectiveness, a the practical results obtained here demonstrate. This analysis can be helpful for future optimizations, enabling the exploration of different configurations of quantum circuits to improve the efficiency of the algorithm in similar contexts ^32,59^.

Studies attempting to generate biological blocks using quantum computing have not yet been reported, and the present study aimed to explore this possibility because quantum computing and synthetic biology are on the rise. Furthermore, the need to quickly and effectively analyze large datasets to accelerate and sensitize the construction of biological building blocks is of great interest in biotechnology, especially in genetic and biological systems engineering.

The discussion presented here highlights and innovative frontier at the intersection of quantum computing and biological sciences, particularly in genomics and genetic engineering. As researchers increasingly explore the applications of quantum algorithms to solve complex biological problems, it becomes clear that significant advances are possible, thereby addressing long-standing challenges in these fields. First, the potential of quantum computing to improve DNA sequence analysis and protein design algorithms cannot be overstated. The ability to leverage quantum superposition and interference offers a paradigm shift in terms of how we process biological data. Traditional computational methods often struggle with the exponential complexity of genomic data, making them unsuitable for large-scale analyses. Owing to its inherent ability to process vast datasets simultaneously, quantum computing offers a viable solution to these challenges.

Furthermore, findings from recent studies highlight the feasibility of achieving substantial speedups in genetic analyses through quantum computing ^13,40^. The potential for quadratic and polynomial speedups over classical methods suggests that quantum technologies could significantly reduce the time required for genome sequencing and variant detection. This speedup is particularly crucial as the volume of genomic data continues to grow exponentially, driven by advances in sequencing technologies and the growing interest in personalized medicine.

Here, our study contributes to the small-scale assessment of the feasibility of accelerating applied analyses with real genomic data and to advancing quantum computing in biological sciences. It is a way to not only keep up with the current hype but also to follow the evolution of a science that, despite being an emerging field, is extremely important for the large volume of biological data that will need to be processed in the future.

In addition, integrating quantum computing with artificial intelligence represents an attractive avenue for future research. By optimizing genetic algorithms and enabling real-time data analysis, quantum computing can revolutionize trait discovery in agriculture and biotechnology. The potential to accelerate the identification of beneficial traits and increase crop resilience can yield significant benefits for food security and sustainability. As the world faces urgent challenges such as climate change and population growth, the ability to innovate agricultural practices through quantum-enabled solutions has become increasingly vital.

Despite these promising advances, it is essential to recognize the challenges that remain regarding the practical implementation of quantum computing in real-world biological applications. The current state of quantum hardware is still evolving, and limitations in qubit availability and coherence time represent significant obstacles. As researchers work to develop more robust and scalable quantum systems, the path to practical applications in biology and genetic engineering will become clearer.

In conclusion, this research represents a significant contribution to the field of applying quantum computing to biology and genetic engineering. The innovative approaches discussed herein provide a foundation for further exploration and experimentation, illuminating the potential of quantum technologies to transform our understanding of complex biological systems. As we advance in this exciting domain, we continue to foster interdisciplinary collaboration, to bridge the gap between quantum computing and the biological sciences. Such synergy will undoubtedly lead to ground-breaking discoveries and innovations, reshaping the biotechnology landscape and offering new solutions to some of the most pressing challenges in healthcare, agriculture, and beyond. The future of quantum computing in biology is auspicious, and continued exploration of its capabilities will be the key to unlocking new frontiers in science and technology.

## Methods

The workflows (Fig. 1) provide an overview of the quantum codes developed in-house and designed for genetic sequence analysis, based on Grover’s Algorithm, on a quantum computing cloud platform. This algorithm is well-suited for bioinformatics applications, specifically for sequence searching within large databases, because it provides a quadratic speedup over classical search algorithms, enabling the handling of the immense scale of genomic data. While classical search methods require O(N) comparisons, where N is the size of the database, Grover’s Algorithm reduces this to O(√N) ^54^. The process begins with two main inputs: the DNA target sequence and the database sequence, in which the superposition of quantum states enables multiple comparisons to co-occur. The DNA target sequence is the specific genetic sequence of interest that needs to be analyzed. This could be a specific gene or segment associated with a particular trait. On the other hand, the database sequence represents a collection of genetic sequences stored in a database. These sequences may come from various sources, such as whole genomes, gene libraries, or databases of genetic variations ^11^.

Once the data is prepared, the core of the process involves applying Grover’s Algorithm, a quantum algorithm designed for efficiently searching unsorted databases. By leveraging quantum computing, Grover’s Algorithm significantly speeds up the search for the target sequence within the database compared to classical methods, reducing the number of comparisons required. The execution of Qubio.py takes place on a quantum computing cloud platform.

After the algorithm runs, the analysis results are obtained. These results indicate the presence or location of the target sequence within the database and are then processed to extract meaningful insights. In scientific research, understanding and analyzing genetic sequences can lead to insights into biological mechanisms, the study of genetic diseases, and the development of new therapies and bioactive molecules. In agriculture, genetic analysis can help identify genes responsible for desirable plant traits, such as disease resistance, productivity, and product quality. Additionally, genetic analysis can be used in environmental monitoring to track biodiversity and assess ecosystem health by detecting and monitoring species and genetic variations.

This code implements Grover’s algorithm to efficiently search for a specific target sequence within a database of DNA sequences. By leveraging Grover’s transformation, represented mathematically by matrices, the algorithm amplifies the probability of locating the desired sequence.

### Sequences definition

To define the short-length target sequence (50 bp), a protein sequence isolated from the Brazilian biomes metagenome project^50^ was explored and analyzed for its functional domain. Its most conserved portion was highlighted and filtered to serve as a search target (fig. 7). The chosen protein, a Cas9-like nuclease, is involved in the CRISPR/Cas9 system, which offers several advantages, including high editing efficiency, ease of operation, cost-effectiveness, and a diverse set of recognition sites. It can be used to edit the human genome to treat genetic defects or illnesses caused by gene mutations. It can also edit plant genes to create more resilient plants. Furthermore, the CRISPR/Cas9 editing technology enables the elimination of pathogens such as bacteria and viruses, potentially curing infectious diseases^60^.

**Fig. 7.**
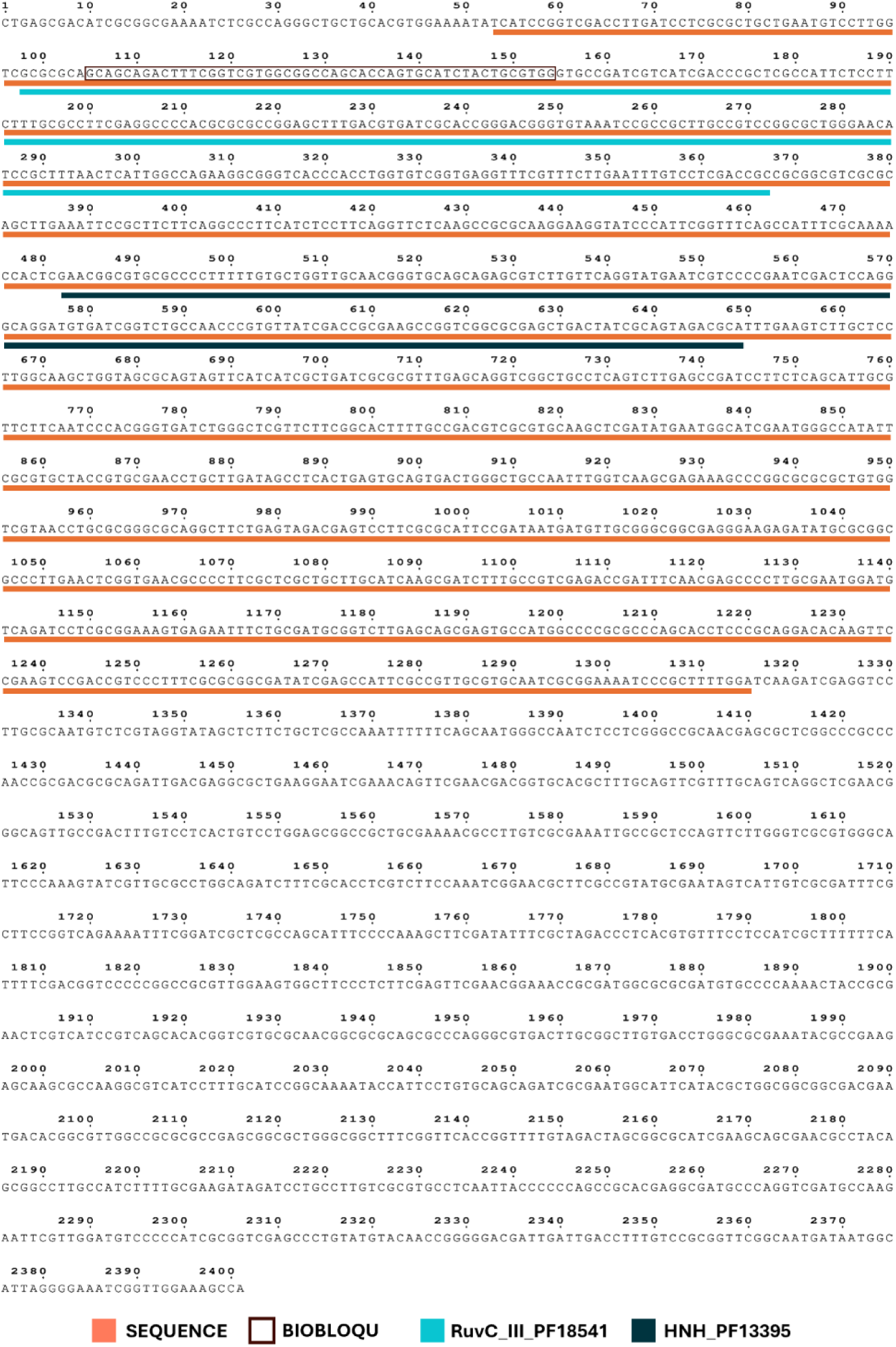
Rational design of the target sequence. Visual identification and rational selection of the conserved target sequence within the RuvC_III domain of Cas9. The orange segments represent the selected target sequence embedded in the Cas9 coding region. The cyan segments correspond to the RuvC functional domain of Cas9, while the dark-blue segments denote the HNH functional domain. The BioBloQu structure was designed based on the conserved region of the RuvC domain, which is encompassed by the defined target sequence.

### Qubio.py code

In this hybrid classical–quantum framework, Grover’s algorithm is employed as the quantum search component. In the Grover circuit, each qubit is initialized through a Hadamard transformation, generating a uniform superposition of |0⟩ and |1⟩. This enables the quantum state to coherently explore all candidate solutions in parallel, a key mechanism underlying quantum amplification. Each Grover iteration increases the probability amplitude associated with the marked state, thereby enhancing the likelihood of identifying the correct subsequence beyond what is achievable with classical search alone.

Before the quantum stage, our pipeline performs classical filtering using the Hamming distance to reduce the search space. This step admits sequences whose dissimilarity to the query is up to approximately 30%, corresponding to about 70% nucleotide similarity between the target sequence and the genomic region under analysis. In practical terms, for a sequence of length 100, this threshold implies that roughly 70 nucleotides are identical while up to 30 may differ. While this level of similarity is often sufficient to infer potential homology, more detailed bioinformatic validation may be required depending on the genomic context.

### Quantum-based hybrid search for DNA fragments

The methodology integrates hybrid classical–quantum code that integrates Hamming-distance filtering with a simulated Grover search applied to a reduced candidate set. The procedure combines theoretical foundations, classical preprocessing, quantum circuit construction, and analysis of oscillatory probability behavior characteristic of Grover’s amplitude amplification.

### Classical filtering and Hamming similarity

The classical stage begins by quantifying similarity between a target DNA fragment and subsequences of a larger reference sequence. This is achieved through the Hamming distance, defined for two sequences a and b of equal length L as:

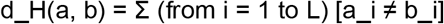

where a_i and b_i represent the nucleotides at corresponding positions, and the indicator [a_i ≠ b_i] equals 1 when the nucleotides differ and 0 otherwise. A sliding window of length L is applied to the reference sequence D, generating segments:

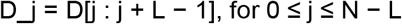

The Hamming distance d_H(T, D_j) is calculated for each window, and the K subsequences with the smallest distances are selected as candidates. The ordered classical candidate set is given by:

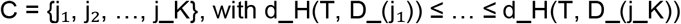

This filtered set forms the reduced search domain to be processed by the quantum routine.

### Grover Algorithm implementation

In the quantum stage, Grover’s algorithm is used to identify the correct candidate among the K classical indices. The algorithm begins with the uniform superposition over all candidates:

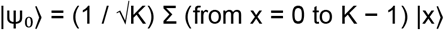

Each Grover iteration consists of two operations: the oracle and the diffusion operator. The diffusion operator performs inversion about the mean and is defined as:

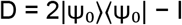

The state after r Grover iterations is:

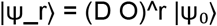

Grover’s amplitude amplification leads to periodic oscillations in the probability of the marked state. The probability of measuring the correct index after r iterations is:

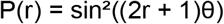

where the angle θ satisfies:

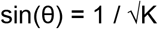

The optimal number of iterations that maximizes the success probability is:

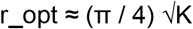

### Oracle Function integration

In the quantum stage, the algorithm includes an explicit oracle function implemented directly in the circuit. This oracle performs a conditional phase inversion on the marked state associated with the target DNA fragment.

In our implementation, the oracle operator follows exactly the standard qubit-based construction used in Grover’s algorithm. The Hilbert space for a target sequence of length *L*is the qubit tensor-product space

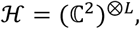

and not ℂ^*L*^or any single-register space. Each nucleotide is encoded as a single-qubit computational-basis state (or superposition) within this tensor structure.

The oracle is a phase-flip reflection acting on the marked basis state ∣ *x*^∗^⟩corresponding to the target DNA fragment, defined as

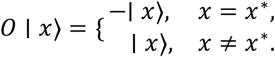

Operationally, the circuit implements this reflection by applying X gates to the qubits that must be inverted to align ∣ *x*^∗^⟩with ∣ 11 … 1⟩, applying a multi-controlled X (MCX) gate to flip the global phase of this basis vector, and then undoing the alignment. The sequence

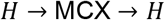

on the final qubit performs the standard Grover conditional phase inversion. The oracle in Qubio.py code is mathematically identical to the canonical Grover oracle acting in the qubit tensor space (ℂ^2^)^⊗*L*^, ensuring that the amplitude amplification dynamics follow the conventional Grover model.

### Data preparation

The experimental process begins by specifying the target DNA sequence T of length L and the reference sequence D of length N. A sliding-window Hamming-distance computation is performed, and the top K subsequences with the smallest distances are selected. Typical experimental choices include L ≈ 50 nucleotides and K equal to 16 or 20.

### Preparation of quantum simulation

To simulate the quantum search, the number of qubits is derived from the size of the candidate set:

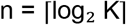

The classical index corresponding to the correct candidate is encoded in an n-bit binary string. A quantum circuit is initialized, and Hadamard gates are applied to all qubits to generate the uniform superposition state |ψ_0_⟩. Grover iterations are then performed, each composed of the oracle marking the correct index and the diffusion operator amplifying its amplitude. Measurements are conducted after r iterations, and the experiment is repeated for r = 1, 2, …, r_max to observe oscillatory behavior. For each iteration count, measurement results are collected to estimate the empirical probability distribution.

### Execution Environment

The quantum execution is performed using the Qiskit Aer simulator to obtain stable probability estimates. The circuit undergoes transpilation prior to execution to optimize its structure for the simulation backend.

### Measure qubits

We implemented Grover’s algorithm on real quantum hardware and compared the outcomes with classical simulations. The quantum simulations were performed using the Qiskit framework, leveraging the Qiskit Aer simulator for up to 27 qubits.

Despite these constraints, the quantum implementation demonstrated the algorithm’s ability to efficiently identify target sequences with far fewer comparisons than classical algorithms. By running the simulations via Google Colab, hosting the Qubio.py code, we highlighted the potential advantages of quantum-based algorithms in scaling up computational tasks, even in bioinformatics applications.

### Scar and BioBloQu code

To search for scar sequences and insert genetic blocks using quantum computing begins with defining the target sequences and the database sequence. Analyzes whether there is a scar, two predefined sequences (left end: AAAATCTGTCATAAATTATC and right end: ATTATTCTCCTTTCTTTAGT) from a specific essential gene in a determined region of the minimal genome of *M. mycoides* JCVI-Syn3B (3906 nucleotides). Part of the minimal cell genome was used as database for search.

The insertion of the genetic block is accomplished by identifying the position of the scar in the database sequence and replacing it with the BioBloQu. The circuit is then simulated using the Qiskit backend, where measuring the qubits results in counts that represent the probability of finding the target sequences.

All codes were run in the IBM simulation environment using jupyter notebook and google colab.

### Quantum encoding of the BioBloQu construct

We developed a quantum encoding framework to represent the full BioBloQu genetic construct as a quantum state, enabling simulation of hybrid biomolecular structures within a digital quantum environment. The BioBloQu unit is composed of four predefined genetic elements—a promoter, a ribosome binding site (RBS), an enzymatic domain of fixed length, and a transcription terminator—which are concatenated to form a single linear sequence:

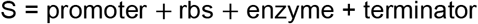

For a sequence S of length L, a quantum register of L qubits is initialized. Each nucleotide is mapped to a quantum operation that prepares its corresponding basis or superposition state. Specifically, the encoding employs base-selective transformations:

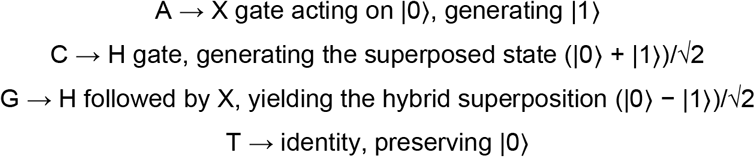

Thus, the encoded BioBloQu state is the tensor product:

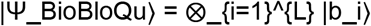

where each |b_i⟩ is the quantum state corresponding to nucleotide S_i_ under the mapping above.

After construction, the full register is measured on a computational basis to characterize the distribution associated with the encoded sequence. The quantum simulation is performed using the Qiskit Aer simulator with 1024 measurement shots, producing a frequency histogram of the observed bitstrings. This encoding stage does not perform amplitude amplification or search; instead, it provides a quantum-representational substrate for synthetic constructs, enabling downstream hybrid workflows that integrate quantum state preparation with classical genome engineering tasks.

This procedure establishes a reproducible quantum representation of a genetic payload and serves as a digital analogue of synthetic DNA assembly in a fully quantum-simulated environment.

### Grover-Based quantum search for scar identification and BioBloQu insertion

To locate genomic scar sites and guide synthetic insertion events, we developed a quantum-assisted search routine based on a Grover amplitude-amplification model implemented on top of classical genome screening. This methodology maps target DNA flanks into quantum states and executes a Grover search over a structured search space derived from the reference genome.

### Classical scar detection

Given a target DNA fragment T of length L and a reference genome sequence D, the algorithm first identifies candidate scar positions using a classical substring match. If T occurs in D, the index j corresponding to the right boundary of the scar is obtained by:

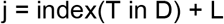

This position defines the insertion coordinate for the BioBloQu construct. The target sequence T also serves as the pattern for the oracle in the quantum stage described below.

### Quantum Oracle construction

For a target sequence T = t_1_t_2_…t_L, a quantum register of L qubits is initialized in uniform superposition through application of Hadamard gates:

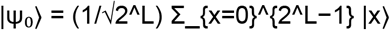

The oracle O acts by flipping the phase of basis states corresponding to the classical target sequence. Operationally, each qubit position encoding an adenine triggers an X gate to align the computational basis with |11…1⟩ before applying a multi-controlled X (MCX) operation, followed by the inverse of the pre-oracle alignment. The oracle therefore implements:

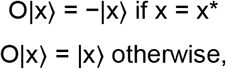

where x* denotes the bitstring encoding the target sequence T.

### Diffusion Operator (amplitude amplification)

The amplification step applies to the Grover diffusion operator:

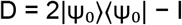

implemented via the standard circuit decomposition:

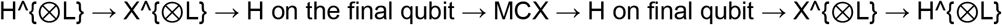

This transformation performs inversion about the mean, increasing the amplitude associated with the marked state.

### Iterative Grover dynamics

A sequence of r Grover iterations is applied, each consisting of oracle followed by diffusion. For a search space of size N = 2^L, the optimal iteration count is:

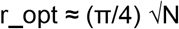

In practice, the implementation approximates this value using:

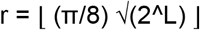

After r iterations, the state evolves to:

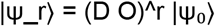

The probability of measuring the correct sequence is given by the Grover oscillation formula:

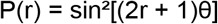

with sin(θ) = 1 / √N.

### Measurement and Probability Estimation

The final quantum state is measured using 8192 shots on the Qiskit Aer simulator. The empirical success probability is:

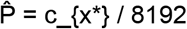

where c_{x*} is the number of occurrences of the marked bitstring.

### BioBloQu insertion into the scar site

Upon successful quantum identification of the target flank, the BioBloQu construct is inserted into the genome at the scar location, yielding:

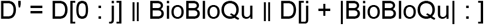

This hybrid strategy couples quantum search with classical genome rewriting, enabling precise synthetic modification driven by quantum-determined sequence identification.

## Code-Data availability

The raw data and code generated and analyzed in this study are available from the corresponding author on request.

## Additional applications

While our study applies Grover’s algorithm to bioinformatics, the quantum search algorithm can be adapted to broader scientific challenges. For example, it could be used to identify patterns in astrobiology datasets or search for signals in massive astronomical data. As quantum computing technology matures, Grover’s algorithm has the potential to overcome computational limitations previously faced by classical systems.

Quantum search can also enable the identification of novel traits within genetic sequences that were previously inaccessible due to computational resource constraints. This could revolutionize fields such as genomics, pharmaceuticals, agriculture, and environmental science, driving forward innovation across a variety of disciplines. Grover’s algorithm offers a transformative approach to solving computational search problems, providing quadratic speedup over classical methods. Quantum computing promises to revolutionize domains like computational biology, but significant challenges remain in hardware scalability and the development of domain-specific quantum algorithms. Collaborative efforts between quantum physicists and computational scientists will be essential to overcome and pushing quantum technology beyond the constraints of classical systems.

## Acknowledgments

We acknowledge the National Institute of Science and Technology in Synthetic Biology, National Institute of Science and Technology in Engineering Systems Biology and the Ministry of Agriculture and Livestock.

## Author contributions

Conceptualization: PVP, ER

Methodology: PVP, DB, MHV, ER

Investigation: PVP, DB, LMAT, RNL, MAO, MHV, ER

Visualization: PVP, DB, MHV, ER

Funding acquisition: ER

Project administration: ER

Supervision: ER

Writing – original draft: PVP, DB, MHV, ER

Writing – review & editing: DB, LMAT, RNL

## Data and materials availability

All the data presented in this paper were generated by numerical simulations using computer programs. Correspondence and requests for materials should be addressed to Elibio Rech (e-mail address). The code underlying this study is not publicly available but may be made available to qualified researchers upon reasonable request from the corresponding author.

## Supplementary Materials

### Code-Data availability

(**The availability of the codes below is provided here as an attachment, exclusively for the editor and reviewers)**.

~~~
QuBio.py - code to search the target_sequence in database_sequence using quantum computing
#install qiskit and qiskit-aer before run on google colab and use IBM quantum simulator
pip install qiskit
pip install qiskit-aer
# ==========================================================
# QUBIO
# ==========================================================
from qiskit import QuantumCircuit, transpile
from qiskit_aer import Aer
from qiskit.circuit.library
import MCXGate
import matplotlib.pyplot as plt
import numpy as np import math
# ==========================================================
# 1. Dataset Preparation #
# The target sequence represents a fragment of interest,
# while the database sequence corresponds to a long DNA chain
# where the algorithm attempts to find the closest match.
# ==========================================================
target_sequence = ‘GCAGCAGACTTTCGGTCGTGGCGGCCAGCACCAGTGCATCTACTGCGTGG’
database_sequence = (
 ‘CTGAGCGACATCGCGGCGAAAATCTCGCCAGGGCTGCTGCACGTGGAAAATATCATCCGGTCGA CCTTGATCCTCGCGCTGCTGAATGTCCTTGGTCGCGCGCAGCAGCAGACTTTCGGTCGTGGCGGCC AGCACCAGTGCATCTACTGCGTGGGTGCCGATCGTCATCGACCCGCTCGCCATTCTCCTTCTTTGCG CCTTCGAGGCCCCACGCGCGCCGGAGCTTTGACGTGATCGCACCGGGACGGGTGTAAATCCGCCG CTTGCCGTCCGGCGCTGGGAACATCCGCTTTAACTCATTGGCCAGAAGGCGGGTCACCCACCTGGT GTCGGTGAGGTTTCGTTTCTTGAATTTGTCCTCGACCGCCGCGGCGTCGCGCAGCTTGAAATTCCG CTTCTTCAGGCCCTTCATCTCCTTCAGGTTCTCAAGCCGCGCAAGGAAGGTATCCCATTCGGTTTCA GCCATTTCGCAAAACCACTCGAACGGCGTGCGCCCCTTTTTGTGCTGGTTGCAACGGGTGCAGCAG AGCGTCTTGTTCAGGTATGAATCGTCCCCGAATCGACTCCAGGGCAGGATGTGATCGGTCTGCCAA CCCGTGTTATCGACCGCGAAGCCGGTCGGCGCGAGCTGACTATCGCAGTAGACGCATTTGAAGTCT TGCTCCTTGGCAAGCTGGTAGCGCAGTAGTTCATCATCGCTGATCGCGCGTTTGAGCAGGTCGGCT GCCTCAGTCTTGAGCCGATCCTTCTCAGCATTGCGTTCTTCAATCCCACGGGTGATCTGGGCTCGTT CTTCGGCACTTTTGCCGACGTCGCGTGCAAGCTCGATATGAATGGCATCGAATGGGCCATATTCGC GTGCTACCGTGCGAACCTGCTTGATAGCCTCACTGAGTGCAGTGACTGGGCTGCCAATTTGGTCAA GCGAGAAAGCCCGGCGCGCGCTGTGGTCGTAACCTGCGCGGGCGCAGGCTTCTGAGTAGACGAGT CCTTCGCGCATTCCGATAATGATGTTGCGGGCGGCGAGGGAAGAGATATGCGCGGCGCCCTTGAAC TCGGTGAACGCCCCTTCGCTCGCTGCTTGCATCAAGCGATCTTTGCCGTCGAGACCGATTTCAACGA GCCCCTTGCGAATGGATGTCAGATCCTCGCGGAAAGTGAGAATTTCTGCGATGCGGTCTTGAGCAG CGAGTGCCATGGCCCCGCGCCCAGCACCTCCCGCAGGACACAAGTTCCGAAGTCCGACCGTCCCT TTCGCGCGGCGATATCGAGCCATTCGCCGTTGCGTGCAATCGCGGAAAATCCCGCTTTTGGATCAA GATCGAGGTCCTTGCGCAATGTCTCGTAGGTATAGCTCTTCTGCTCGCCAAATTTTTTCAGCAATGG GCCAATCTCCTCGGGCCGCAACGAGCGCTCGGCCCGCCCAACCGCGACGCGCAGATTGACGAGGC GCTGAAGGAATCGAAACAGTTCGAACGACGGTGCACGCTTTGCAGTTCGTTTGCAGTCAGGCTCGA ACGGGCAGTTGCCGACTTTGTCCTCACTGTCCTGGAGCGGCCGCTGCGAAAACGCCTTGTCGCGAA ATTGCCGCTCCAGTTCTTGGGTCGCGTGGGCATTCCCAAAGTATCGTTGCGCCTGGCAGATCTTTCG CACCTCGTCTTCCAAATCGGAACGCTTCGCCGTATGCGAATAGTCATTGTCGCGATTTCGCTTCCGG TCAGAAAATTTCGGATCGCTCGCCAGCATTTCCCCAAAGCTTCGATATTTCGCTAGACCCTCACGTGT TTCCTCCATCGCTTTTTTCATTTTCGACGGTCCCCCGGCCGCGTTGGAAGTGGCTTCCCTCTTCGAG TTCGAACGGAAACCGCGATGGCGCGCGATGTGCCCCAAAACTACCGCGAACTCGTCATCCGTCAGC ACACGGTCGTGCGCAACGGCGCGCAGCGCCCAGGGCGTGACTTGCGGCTTGTGACCTGGGCGCG AAATACGCCGAAGAGCAAGCGCCAAGGCGTCATCCTTTGCATCCGGCAAAATACCATTCCTGTGCAG CAGATCGCGAATGGCATTCATACGCTGGCGGCGGCGACGAATGACACGGCGTTGGCCGCGCGCCG AGCGGCGCTGGGCGGCTTTCGGTTCACCGGTTTTGTAGACTAGCGGCGCATCGAAGCAGCGAACG CCTACAGCGGCCTTGCCATCTTTTGCGAAGATAGATCCTGCCTTGTCGCGTGCCTCAATTACCCCCC AGCCGCACGAGGCGATGCCCAGGTCGATGCCAAGAATTCGTTGGATGTCCCCCATCGCGGTCGAG CCCTGTATGTACAACCGGGGGACGATTGATTGACCTTTGTCCGCGGTTCGGCAATGATAATGGCATT AGGGGAAATCGGTTGGAAAGCCACGGCAAAGTGGTTTTACCACTGAAGGATCACCCCCGTTCGGCG ACCCTGAACGGGGGGATTCCTTAATATCAACTGTCATTACGGCGGGCTCACATTCGCTCACTCAGAA AGCTGGATCTGTCAGCTTTGTTCGTTATTAAAGGCATGCGGCTTGCGCCCTTGACAAAAGCACGTTG TCGCCTTCGTTACGGTCGAACGAAGCTTGCGACCGTGAGTTCCGGCA’
)
# ==========================================================
# 2. Classical Preprocessing Functions #
# These routines emulate biological sequence alignment logic
# (Hamming distance sliding window) to select a small number
# of best candidate fragments for quantum evaluation.
# ==========================================================
def hamming(a, b):
 “““Compute Hamming distance between two strings.”““
 return sum(ch1 != ch2 for ch1, ch2 in zip(a, b))
def find_candidates(target, database, top_k=16):
  “““
  Classical sliding window that returns the top_k database
  fragments with the smallest Hamming distance from the target.
  “““
L = len(target)
scores = []
for start in range(len(database) - L + 1):
  frag = database[start:start+L]
  score = hamming(target, frag)
  scores.append((score, start, frag))
scores.sort(key=lambda x: x[0])
return scores[:top_k]
# ==========================================================
# 3. Quantum Core: Grover Search on Candidate Indices #
# A reduced version of Grover’s algorithm applied to the
# discrete set of top-K candidate positions in the database.
# ==========================================================
def int_to_bitstr(x, n):
   return format(x, f’0{n}b’)
def mark_index_oracle(qc, index_bits):
“““Oracle: phase-inverts the state corresponding to index_bits.”““
n = len(index_bits)
for i, b in enumerate(index_bits):
  if b == ‘0’:
   qc.x(i)
qc.h(n - 1)
if n > 1:
  qc.append(MCXGate(n - 1), list(range(n - 1)) + [n - 1])
else:
  qc.z(0)
qc.h(n - 1)
for i, b in enumerate(index_bits):
  if b == ‘0’:
   qc.x(i)
def diffuser_idx(qc, n):
“““Standard Grover diffusion operator.”““
qc.h(range(n))
qc.x(range(n))
qc.h(n - 1)
qc.append(MCXGate(n - 1), list(range(n - 1)) + [n - 1]) qc.h(n - 1)
qc.x(range(n))
qc.h(range(n))
def grover_on_candidates(best_idx, K, max_iter=10):
“““
Runs Grover’s algorithm over K candidate indices.
Returns a list of measured probabilities for the target index.
“““
n = math.ceil(math.log2(K))
backend = Aer.get_backend(‘qasm_simulator’)
probs = []
target_bits = int_to_bitstr(best_idx, n)
for k in range(1, max_iter + 1):
  qc = QuantumCircuit(n)
  qc.h(range(n))
  for _ in range(k):
    mark_index_oracle(qc, target_bits)
    diffuser_idx(qc, n)
 qc.measure_all()
 result = backend.run(transpile(qc, backend), shots=2048).result()
 counts = result.get_counts()
 total = sum(counts.values())
 prob = counts.get(target_bits[::-1], 0) / total # Qiskit uses little-endian
 probs.append(prob)
return probs
# ==========================================================
# 4. Experiment Workflow #
# Step 1: Classical filtering of candidate subsequences.
# Step 2: Quantum search over indices of best candidates.
# Step 3: Probability visualization.
# ==========================================================
TOP_K = 20
MAX_ITER = 10
candidates = find_candidates(target_sequence, database_sequence, top_k=TOP_K)
if not candidates:
  raise SystemExit(“No candidate fragments found.”)
best_score, best_start, best_frag = candidates[0]
print(f”Best candidate (start={best_start}) with {best_score} mismatches.”)
K = len(candidates)
best_idx = 0 # since list is sorted by best score
probs = grover_on_candidates(best_idx, K, max_iter=MAX_ITER)
# ==========================================================
# 5. Visualization and Analysis #
# The probability curve shows the oscillatory nature of
# Grover’s amplification, with a distinct optimal iteration
# maximizing the chance of measuring the best candidate.
# ==========================================================
plt.figure(figsize=(8, 5))
plt.plot(range(1, MAX_ITER + 1), probs, marker=‘o’)
plt.xlabel(‘Number of Grover iterations’)
plt.ylabel(‘Probability of measuring the best candidate’)
plt.title(f’Qubio over candidate indices (K={K}, best start={best_start}, mismatches={best_score})’)
plt.grid(True, linestyle=‘--’, alpha=0.6)
plt.ylim(0, 1) plt.show()
for i, p in enumerate(probs, 1):
  print(f”Iteration {i:2d}: prob = {p:.3f}”)
##Scar identification and BioBloQu addition code
from qiskit import QuantumCircuit, transpile
from qiskit_aer import Aer
import matplotlib.pyplot as plt
import numpy as np
# ----------------------------------------------------------------
# Defining the target sequences (right and left scar flanks)
# and the reference database genome sequence (JCVI-Syn3B)
# ----------------------------------------------------------------
target_sequences = [‘AAAATCTGTCATAAATTATC’, ‘ATTATTCTCCTTTCTTTAGT’]
database_sequence = (
‘ATTTTTTTCTTTCTAAATACTTTTATATTTTATTTTAAATTCTTATGAATTTGATTTAAATAAGTCTGTTTA TTATTAATATCTTCAATATAAGCTGAAATTAAAATTCTAATAACTGGATCATTTATTTTTTTATTAATAGC TTTAATTATGTTAAATAAGTTTTTTTGAAAGTCTAATTCTAATCTTTCAAATTGATAGTTAAACATTAAAT TATTATCATAAGCTTTATTAAATTCAATCCAATGATTATTTAAAGCATCATTAAAAATTCAATTAATAAAA ATTTGATTAATAAAATCTCTTAATAATGTATAACTAGTTAATAAAAACTTTTTATAATTAATTTCAATAAA ATTAGTGTTATTTGTTTTTAATAAATAATCAATACAAGCAGTATTTGTTTTTTTAAATAAATTTCTAATTT GATTAATTTTATTTTCAATAGCATCAATCATTAAATAAGAAAAGTTTAATTTAGATGAATTTTGTAAATAT GTAATGAAATTAGGTTTATTTAAAATATCTTCAAAATCAGAAAAACTTCAATTAATAGTCATATTTAAAT ATTTAGTTTTTTGAATTTTTGAAACATTATTTTTAAAAAAACTATTTCTAATTAATATAAAATCTGTCATA AATTATCATTATTCTCCTTTCTTTAGTCTATTATTATTATAAATTAACATAATATTTTATATAAATTATTAA TAAAAGAAATAATTTAAAAATAAAAAAAGAACCACTTAAGGTTCTTTAGAAACTGGTAGCTTGCTATCT TCTCACATAAGTAATATTTTCACCGTAATAGAGCTTAACTTCTGTGTTCGGCATGGGAACAGGTGTGA CCTCTATGCTATAACCACCAGATTATTTTTTGATTGTTCTTTGAAAACTGATTATTAGCTTGAAATGTAA GTTAAAATTGCATTTTCTAAAATTCACTCGATCTATTAGTAATGGTTAGCTTAATGCCTCACAGCACTT ACACATCCATCCTATCAACCATGTGGTCTACATGGGATCTTACATCTAAAAGAATGGGAAAATTCATC TTAAAGGAGGCTTCTCGCTTAGATGCCTTCAGCGATTATCCTTTCTGCACATAGCTACCCTGCTGTGC CACTGGCGTGACAACAGGAGCACCAGGGGTGCATCCATTCCGGTCCTCTCGTACTAGGAACAGCTC TTTTCAATTTTCCAACGCCCACAACAGATAGGGACCAAACTGTCTCACGACGTTCTGAACCCAGCTC GCGTACCGCTTTAATGGGCGAACAGCCCAACCCTTGGAACCGACTACAGCTCCAGGATGCGATGAG CCGACATCGAGGTGCCAAACCTCCCCGTCGATGTGAACTCTTGGGGGAGATAAGCCTGTTATCCCC GGGGTAACTTTTATCCGTTGAGCGACGGCCCTTCCACACGGGACCGCCGGATCACTAAGTCCTACT TTCGTATCTGTTCGACTTGTAAGTCTCGCAGTTAAGCATCCTTCTACCTTTGCGCTCTATGTATGATTT CCAACCATACTGAGGATACCTTTGAGCGCCTCCGTTACTTTTTGGGAGGCGACCGCCCCAGTCAAAC TACCCACCAGACACTGTCCTTAATCCAGATAATGGATCCAAGTTAGAAATCCAATATAACGAGGGTG GTATTCCAAGGTTGACTCCACTAGAACTAGCGTCCTAGTCTCATAGTCTCCCACCTATCCTCTACACG TCATACCAAATTTCAATATCAAGTTATAGTAAAGCTCCACGGGGTCTTTCCGTCTAGTTGCGGGTAAC CGGCATCTTCACCGGTACTAAAATTTCACCGAGTCTGCAGCCGAGACAGCGAAGGGATCATTACACC TTTCGTGCGGGTCAGAATTTACCTGACAAGGAATTTCGCTACCTTAGGACCGTTATAGTTACGGCCG CCGTTCACCGGGGCTTCAATTCAAAGCTTCGCTTGCGCTGACTTCTCCTTTTAACCTTCCGGCACTG GGCAGGTGTCACCCCCTATACATCGTCTTGCGACTTAGCAGAGAGCTGTGTTTTTGTTAAACAGTTG CCCCTCCCTCTTCACTGCGGCTTATCATAGATAAGCACTCCTTCTTCCGAAGTTACGGAGTTATTTTG CAGAGTTCCTTAGCTACAGTTATCTCGCTTGCCTTAGGATTTTCTCCTTGACCACGTGTGTTCGTTCT AGGTACAGGCAGTTAATTATTAAGTTAGAAGCTTTTCTTGGAAGCATGGAGTCATATACTTCGTTACT AGGCGAACCGTTCACTCCCCTATTTTCACACTTCAATGTTAATACAATGCGGATTTGCCAACATTGCC ATCTTTGTGCTTAGCCCGGGACATCCAACACCCGGGATATACTATCCTTCTTCGTCACTCCATCACTA ATTAACTGGTACAGGAATATCAACCTGTTGTCCATCGACTACGCTTTTCAGCCTCGCCTTAGGTCCTG ACTAACCCTGGGTGGACGAACCTTGCCCAGGAAACCTTGGTCAAACGGCATGGAAGATTCTCACTTC CAAACGTTACTCATGCCGGCATAATCACTTCTAAACACTCCACCAGTCTTCACAGTCTGACTTCATCA TATTTAGAACGCTCCCCTACCACTGTATTTACATACAATCCGTAGCTTCGGTGGTAAACTTAAGCCCC GGTACATTTTCGGCGCAGAATCACTCGACTAGTGAGCTGTTACGCACTCTTTAAATGATGGCTGCTT CTAAGCCAACATCCTAGCTGTCTGTGCAGTTCCACATCCTTACACACTTAGTTTACACTTAGGGACCT TAGCTGACGATCTGGGCTGTTTCCCTCACGAGCATGGACCTTATCACCCATGTTCTGACTGCTGTGC ACAAATTATGGCATTCGGAGTTTAATTCTAATCAGTACCGCTAGGTGCAGCCATCATAGATTCAGTGC TCTACCTCCACAACAATTCACACAACGCTATCCTTAAAGATATATCGGGGAGAACTAGCTATCTCCAG GTTCGATTGGAATTTCACCGCTAGCCACAAGTCATCCAAAGTCTTTTCAACGAATACTGGTTCGGTCC TCCATTAGGTTTTACCCTAACTTCAACCTGCTCATGGCTAGATCACCTGGTTTCGTGTCTACTACTGC ATACTAAACGCCCTATTAAGGCTCGCTTTCACTACGGCTCCGTTTATATCAACTTAACCTTGCATGCA ATAGTAACTCGCCGGCTCTTTCTACAAAAAGCACGGTATTACCCATTAACGGGCTCTACCTTCTTGTA AGCATATGGTTTCAGGGACTATTTCACTCCCCTTTCGGGGTGCTTTTCACCTTTCCCTCACGGTACTG GTTCACTATCGGTAAAATGGTAGTATTTAGGCTTACCAAGTGGTCTTGGCAGATTCCGACAAAATTTC ACGTGTTTCGCCGTACTCAGGATACTTTATTGAGGTCATTACATTTCATATACGGGGCTATCACCCTC TATGGCGCTACTTCCCAGTAGCTTCTATTATATAATGACTTTCTAACTCTTAAAAAAGTCCTACAACCC CACTCCGTAAAGTGGTTTGGCCTGTTCCGTTTTCGCTCGCCGCTACTAACAGAATCATTATTATTTTC TTTTCCTCTAGGTACTAAGATGTTTCAGTTCCCTAGGTTCCCTTCTTATAAGCTATGTATTTACTTATA GATAACATGAGATAAATCATGTCAGGTTTCCCCATTCGGAAATTCCGGGATCGAAGCTCACTTCCAG CTCCCCCAGACTTATCGCAGGTAGTCACGTCCTTCTTCGGCTCCATTTTCCAAGGCATTCACCATATG CTCTTACTATTTTTTAGAAAATCTTACTAGCAATTTGTAACTCAATTAATTTTAGTTAATTGTTTTTGATG TCATTAC’
)
# ----------------------------------------------------------------
# Defining the BioBloQu structure components
# ----------------------------------------------------------------
enzyme_sequence =
“ATGCCACGCAGTAGATGCACTGGTGCTGGCCGCCACGACCGAAAGTCTGCTGCTAA” # 50
nucleotides
promoter = “GCCACCATGGTGAGCAAGGGCGAGGCT”
# Example promoter rbs = “AGGAGG” # Ribosome binding site
terminator = “AATAAA” # Transcription terminator
# ----------------------------------------------------------------
# Function to build a Quantum Circuit representing the BioBloQu construct
# ----------------------------------------------------------------
def create_BioBloQu_circuit(enzyme_sequence, promoter, rbs, terminator):
  total_sequence = promoter + rbs + enzyme_sequence + terminator
  num_parts = len(total_sequence)
 qc = QuantumCircuit(num_parts, num_parts)
 # Encoding nucleotides as quantum states
 for i, base in enumerate(total_sequence):
    if base == ‘A’:
       qc.x(i) # Represents |1>
    elif base == ‘C’:
       qc.h(i) # Superposition
    elif base == ‘G’:
       qc.h(i)
       qc.x(i) # |0> + |1> hybridization
     elif base == ‘T’:
       pass # Represents |0>
qc.measure(range(num_parts), range(num_parts))
return qc, total_sequence
# ----------------------------------------------------------------
# Quantum simulation routine
# ----------------------------------------------------------------
def simulate_circuit(qc):
  backend = Aer.get_backend(‘qasm_simulator’)
  job = backend.run(qc, shots=1024)
  result = job.result()
  counts = result.get_counts(qc)
  return counts
# ----------------------------------------------------------------
# Quantum Oracle for target sequence marking
# ----------------------------------------------------------------
def oracle(qc, target_sequence):
   for i, char in enumerate(target_sequence):
     if char == ‘A’:
       qc.x(i)
 qc.h(len(target_sequence) - 1)
 qc.mcx(list(range(len(target_sequence) - 1)), len(target_sequence) - 1)
 qc.h(len(target_sequence) - 1)
 for i, char in enumerate(target_sequence):
  if char == ‘A’:
      qc.x(i)
# ----------------------------------------------------------------
# Diffusion Operator (Amplitude Amplification)
# ----------------------------------------------------------------
def diffusion_operator(qc, num_qubits):
  qc.h(range(num_qubits))
  qc.x(range(num_qubits))
  qc.h(num_qubits - 1)
  qc.mcx(list(range(num_qubits - 1)), num_qubits - 1)
  qc.h(num_qubits - 1)
  qc.x(range(num_qubits))
  qc.h(range(num_qubits))
# ----------------------------------------------------------------
# Quantum-Guided Search and BioBloQu Insertion
# ----------------------------------------------------------------
def search_target_sequences(target_sequences, database_sequence, bio_block):
   results = []
for target_sequence in target_sequences:
  if target_sequence in database_sequence:
  # Identify scar position and define insertion region
  start_idx = database_sequence.index(target_sequence) + len(target_sequence)
  end_idx = start_idx + len(bio_block)
  # Insert the synthetic BioBloQu block into the scar site
  new_sequence = (
   database_sequence[:start_idx]
   + bio_block
   + database_sequence[end_idx:]
)
 # Quantum search configuration
 num_qubits = len(target_sequence)
 qc = QuantumCircuit(num_qubits)
 qc.h(range(num_qubits))
 oracle(qc, target_sequence)
 num_iterations = int(np.floor(np.pi / 8 * np.sqrt(2 ** num_qubits)))
 for _ in range(num_iterations):
   diffusion_operator(qc, num_qubits)
   oracle(qc, target_sequence)
qc.measure_all()
 backend = Aer.get_backend(‘qasm_simulator’)
 job = backend.run(qc, shots=8192)
 result = job.result()
 counts = result.get_counts(qc)
most_frequent_result = max(counts, key=counts.get)
probability = counts[most_frequent_result] / 8192
results.append((probability, target_sequence, new_sequence))
return results
# ----------------------------------------------------------------
# Main execution block
# ----------------------------------------------------------------
BioBloQu_circuit, total_sequence = create_BioBloQu_circuit(enzyme_sequence, promoter, rbs,
terminator)
result_counts = simulate_circuit(BioBloQu_circuit)
results = search_target_sequences(target_sequences, database_sequence, total_sequence)
for probability, target_sequence, new_seq in results:
 print(f”Target sequence identified: {target_sequence}”)
 print(f”Probability of detection: {probability:.2f}”)
 print(f”New sequence after BioBloQu insertion:\n{new_seq}\n”)
# ----------------------------------------------------------------
# Probability Visualization
# ----------------------------------------------------------------
 plt.figure(figsize=(6, 4))
 probabilities = [result[0] for result in results]
 plt.bar(range(1, len(probabilities) + 1), probabilities, color=‘blue’, alpha=0.9)
 plt.xlabel(‘Target Sequences’)
 plt.ylabel(‘Probability of Finding the Scar Sequence’)
 plt.title(‘Quantum Probability of Scar Site Identification’)
 plt.axhline(y=1, color=‘red’, linestyle=‘--’, label=‘Ideal Probability = 1’)
 plt.grid(axis=‘y’)
 plt.xticks(range(1, len(probabilities) + 1, 1))
 plt.legend()
 plt.tight_layout()
 plt.show()
~~~

